# The effect of external lateral stabilization on ankle moment control during steady-state walking

**DOI:** 10.1101/2022.04.29.490037

**Authors:** A.M. van Leeuwen, J.H. van Dieën, S.M. Bruijn

## Abstract

External lateral stabilization can help identify stability control mechanisms during steady-state walking. The degree of step-by-step foot placement control and step width are known to decrease when walking with external lateral stabilization. Here, we investigated the effect of external lateral stabilization on ankle moment control in healthy participants. Ankle moment control complements foot placement, by allowing a corrective center-of-pressure shift once the foot has been placed. This is reflected by a model predicting this center-of-pressure shift based on the preceding foot placement error. Here, the absolute explained variance accounted for by this model decreased when walking with external lateral stabilization. In other words, we found a reduction in the contribution of step-by-step ankle moment control to mediolateral gait stability when externally stabilized. Concurrently, foot placement error and the average center-of-pressure shift remained unchanged.

## Introduction

Walking without falling requires active stabilization of gait in the mediolateral direction (Bauby and Kuo, 2000). The main mechanism to achieve mediolateral stability during steady-state walking is center-of-pressure control (Hof, 2007; van den Bogaart et al., 2021). Generally, the center-of-pressure is controlled through foot placement (Bruijn and van Dieën, 2018; van Leeuwen et al., 2020) with a complementary role of ankle moment modulation after the foot is placed (Hof et al., 2007; van Leeuwen et al., 2021). Both foot placement and ankle moment control are actively driven (Fettrow et al., 2019; Rankin et al., 2014; van Leeuwen et al., 2021; van Leeuwen et al., 2020), and coordinate the center-of-pressure with respect to variations in the center-of-mass kinematic state (Hurt et al., 2010; Wang and Srinivasan, 2014).

Variance in the center-of-mass kinematic state does not fully explain the variance in foot placement (Wang and Srinivasan, 2014). Foot placement errors remain, which appear to be partially corrected for by ankle moments during single stance (Hof et al., 2007; van Leeuwen et al., 2021). Steps that are too medial are followed by lateral center-of-pressure shifts caused by ankle inversion moments, and vice versa steps that are too lateral are followed by medial center-of-pressure shifts caused by ankle eversion moments (van Leeuwen et al., 2021).

Here, we sought to provide further evidence for the role of ankle moment control in mediolateral steady-state gait stability. Although we identified a relationship between foot placement errors and ankle moments (van Leeuwen et al., 2021), our previous work did not demonstrate a direct relationship between stability and this ankle moment control. In the current study, we bridged this gap by asking participants to walk normally and with external lateral stabilization (Mahaki et al., 2019). External stabilization manipulates the stabilizing demands of gait, and thus allows us to assess the relationship between stability and ankle moment control directly. Establishing this link between ankle moment control and steady-state gait stability, helps to better understand how neurologically-intact individuals maintain their stability during steady-state walking. Such knowledge could help design better treatment or training options, for those with impaired steady-state stability control, such as stroke patients (Dean and Kautz, 2015; Stimpson et al., 2019) or older adults (Arvin et al., 2018).

External lateral stabilization reduces the need for active mediolateral stability control. Therefore, mechanisms of which the use decreases whilst walking with external lateral stabilization, can be identified as stability control mechanisms, active during normal steadystate walking. As such, comparable to perturbations being used to assess reactive stability mechanisms, assistive mechanisms such as external lateral stabilization can provide a tool to assess mechanisms stabilizing steady-state gait. With external stabilization, stability improves (Bruijn et al., 2015), whilst step width and the degree of foot placement control decrease (Mahaki et al., 2019), concurrent with decreased cortical control (Bruijn et al., 2015) and vestibular contributions to gait stability (Magnani et al., 2021). This shows, that during normal walking, the wider steps and the higher degree of foot placement control contribute to mediolateral gait stability. Here, we investigated whether the contribution of ankle moment control also decreases during stabilized walking, to assess its contribution to mediolateral gait stability. We hypothesized that the (step-by-step) contribution of ankle moments decreases during stabilized as compared to normal walking.

## Methods

### Participants

Twenty healthy adults participated in the experiment. Our inclusion criterion was that they were able to walk for approximately one hour. Our exclusion criteria were sports injuries, motor impairments affecting their gait pattern and self-reported balance issues. All participants signed informed consent and ethical approval was granted (VCWE-2020-157) by the local ethical committee of the faculty of Behavioral and Movement Sciences at the Vrije Universiteit Amsterdam.

### Protocol

As part of a more extended protocol, participants walked on a dual-belt treadmill (Motek-Forcelink, Amsterdam, Netherlands) in two conditions, i.e. normal walking and stabilized walking. During normal walking, participants could walk on both of the belts, yet when stabilized, the frame’s position constrained them to the right belt only. Each condition lasted ten minutes. Participants performed a familiarization trial until it felt almost natural to walk in the stabilization device or ten minutes had passed. They were instructed not to resist the lateral stabilization.

All trials were performed at 1.25 (m/s) · 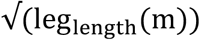 (Hof, 1996). This corresponded to normal walking speed. Participants were instructed fix their gaze at a crosshair, approximately at four meters distance and eye height.

**Table 1.**
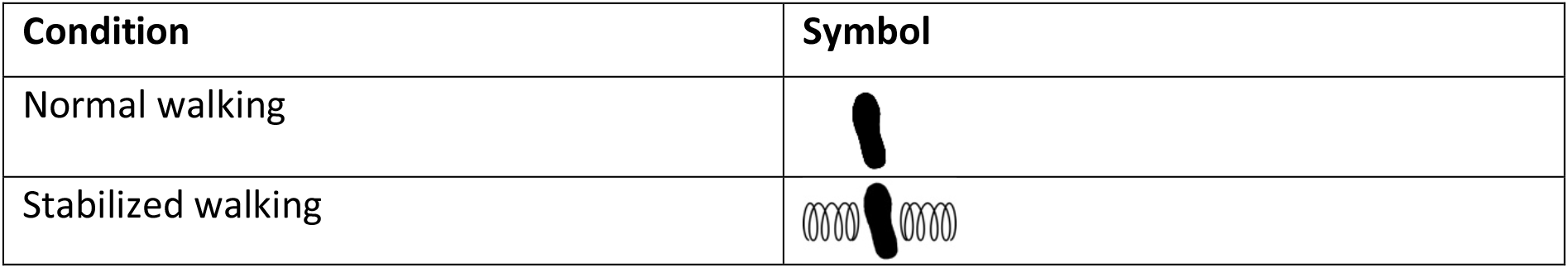
Experimental conditions

| Condition | Symbol |
| --- | --- |
| Normal walking |  |
| Stabilized walking |  |

### Equipment & data collection

#### External lateral stabilization

The springs of the external lateral stabilization pulled the trunk in mediolateral direction at the pelvis level. Sliders allowed for forward translation and rotation around the longitudinal axis (Mahaki et al., 2021). Yet, participants sometimes walked too far forward, beyond the range of motion of the sliders. The experimenter would then ask participants to walk more to the back of the treadmill, unless this disrupted the participant’s performance too much.

#### Kinematics

Cluster markers on the thorax and feet were used to record kinematics using two Optotrak cameras (Northern Digital Inc, Waterloo Ontario, Canada), sampled at 100 Hz. Anatomical landmarks belonging to the thorax and feet segments were digitized in separate recordings, using a six-marker probe.

#### Force plate

Ground reaction forces and moments were recorded from the force plates embedded in the dual-belt treadmill (Motek-Forcelink, Amsterdam, Netherlands), sampled at 200 Hz.

### Data analysis

The code for the analysis can be found on Zenodo https://doi.org/10.5281/zenodo.6994887.)

### Kinematics

The kinematic data were upsampled to 200 Hz (i.e the force plate sampling rate). Digitized anatomical landmarks of the feet (calcaneus and toe tip II) were used to construct local coordinate systems for the feet.

### Gait event detection

Optotrak and force plate data were combined to determine the gait events (heelstrikes and toe-offs). For each trial, the last 400 strides were included.

### Foot placement control

To quantify the degree of foot placement control, and foot placement errors, we considered the foot placement model (Model 1), based on (Wang and Srinivasan, 2014):

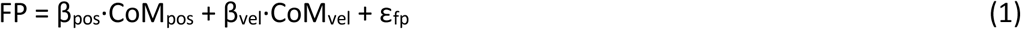

Step width was calculated based on the distance between the calcanei. FP represents the demeaned step widths. CoM_pos_ and CoM_vel_, represent respectively the demeaned center of mass position relative to the stance foot and demeaned center of mass velocity. Jointly these variables describe the center-of-mass kinematic state at terminal swing. ε_fp_ represents the error between actual and predicted foot placement. The estimated thorax center-of-mass was considered as a proxy of the whole-body center-of-mass. The relative explained variance (R^2^) represents the degree of foot placement control. ε_fp_ was used as input for Model 2, to predict the mediolateral center-of-pressure shift based on the preceding error in foot placement.

### Ankle moment control

For each step, CoP displacement was determined during single stance as the average CoP position along the mediolateral axis of the local coordinate system of the foot, with respect to the initial CoP position at contralateral toe-off (Hof and Duysens, 2018). “Average CoP shift” was computed as the average CoP displacement across steps.

As in (van Leeuwen et al., 2021), we considered step-by-step ankle moment control as the relationship between foot placement error (ε_fp_) and the subsequent CoP displacement during single stance (Model 2). CoPshift was computed by demeaning CoP displacement (calculated as described above) and defined such that positive CoP shifts represented a lateral CoP displacement.

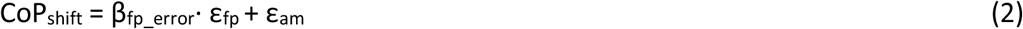

We calculated the absolute variance explained by Model 2 as the contribution of step-by-step ankle moment control, as in (van Leeuwen et al., 2021).

### Gait stability

We computed local divergence exponents (Bruijn et al., 2013; Bruijn, 2021; Mehdizadeh, 2019; Rosenstein et al., 1993). We took the 3D CoM velocity and resampled the signal, so that on average, each stride was 100 samples in length (i.e. we resampled the entire time series of 400 strides to 40.000 data points, retaining potential temporal variations between strides). Then, we created a six-dimensional state space with 25 samples delayed copies. For each timepoint we defined the nearest neighbors as the five points with the smallest Euclidian distances, outside the range of half a stride before or after that timepoint. Subsequently, we tracked the divergence between the timepoint and its nearest neighbors. Using a least-squares-fit, we fitted a line over the first 50 samples of the average logarithmic divergence curve. The slope of this line represents the local divergence exponent.

### Statistics

First, we tested several expectations based on earlier findings (E1-4), (Bruijn et al., 2015; Mahaki et al., 2019; van Leeuwen et al., 2021).

For the normal walking condition, we tested the regression coefficient β_fp_error_ (Model 2) against zero, to replicate our earlier findings (van Leeuwen et al., 2021), that foot placement error significantly predicts the subsequent CoP shift (E1).

We used paired-samples t-tests to assess the effect of stabilization on the degree of foot placement control (R^2^, Model 1), average step width and the local divergence exponent, to confirm earlier findings on walking with external lateral stabilization. We expected a diminished degree of foot placement control (E2), a decreased step width (E3) and a lower local divergence exponent (E4).

To test our hypothesis, we performed paired-samples t-tests for our two main outcome measures, the step-by-step contribution of ankle moment control (absolute explained variance Model 2) and the mean CoP shift (i.e. the overall contribution of ankle moment control) of both conditions.

Finally, we checked whether changes in foot placement error confounded the effect of lateral stabilization, by comparing foot placement precision (the standard deviation of ε_fp_) between normal and stabilized walking.

We treated *p<0.05* as significant.

## Results

Foot placement error (ε_fp_, Model 2) predicted the subsequent CoP displacement during subsequent single stance (CoP_shift_, Model 2), i.e. the regression coefficient β_fp_error_ was significantly different from zero (*p<0.001*). β_fp_error_ was negative, indicating that a too medial/too lateral step was corrected for by a CoP shift in the opposite direction (Figure 2).

**Figure 1.**
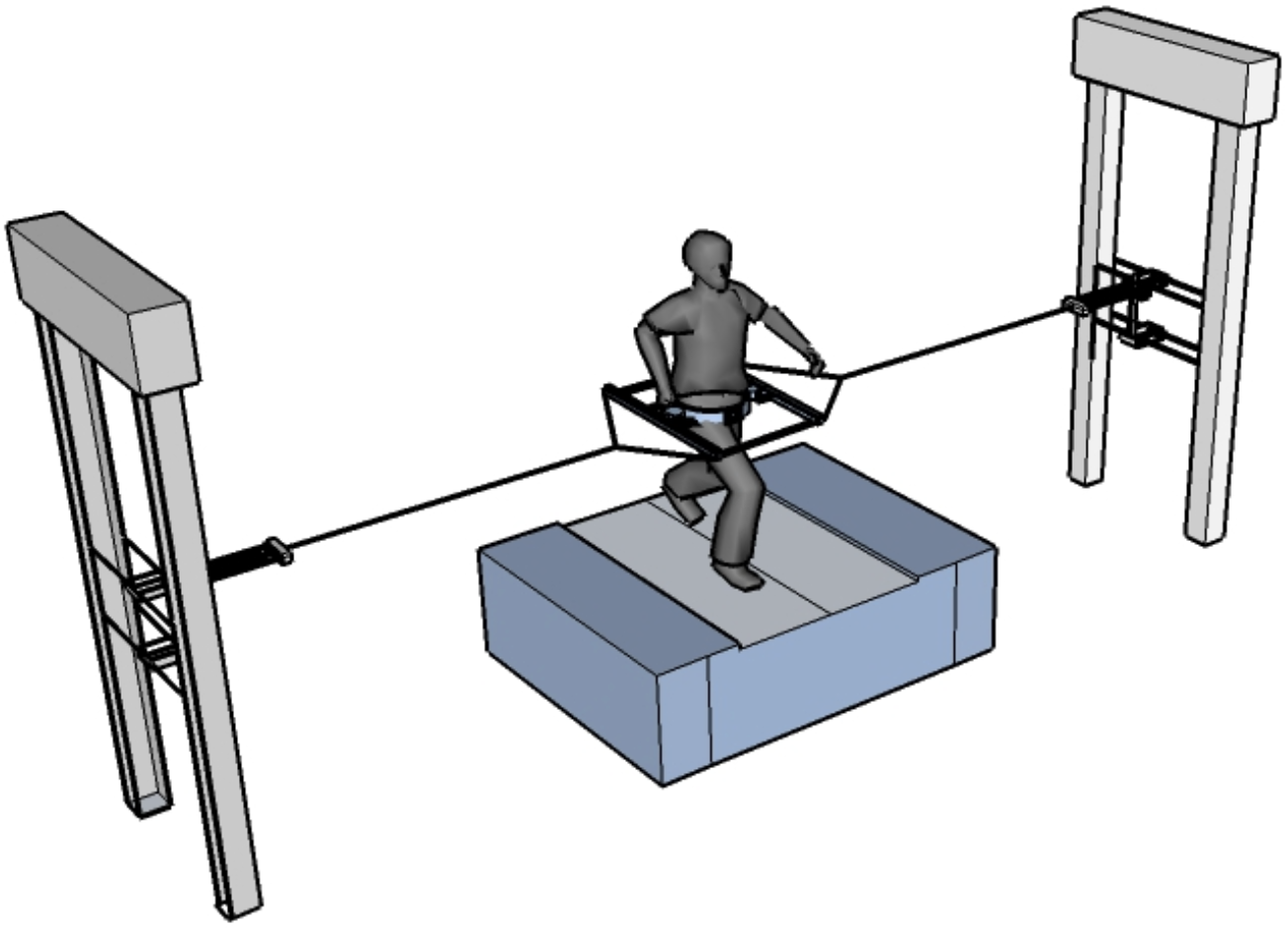
Treadmill with external lateral stabilization. Figure from (Mahaki et al., 2019).

**Figure 2.**
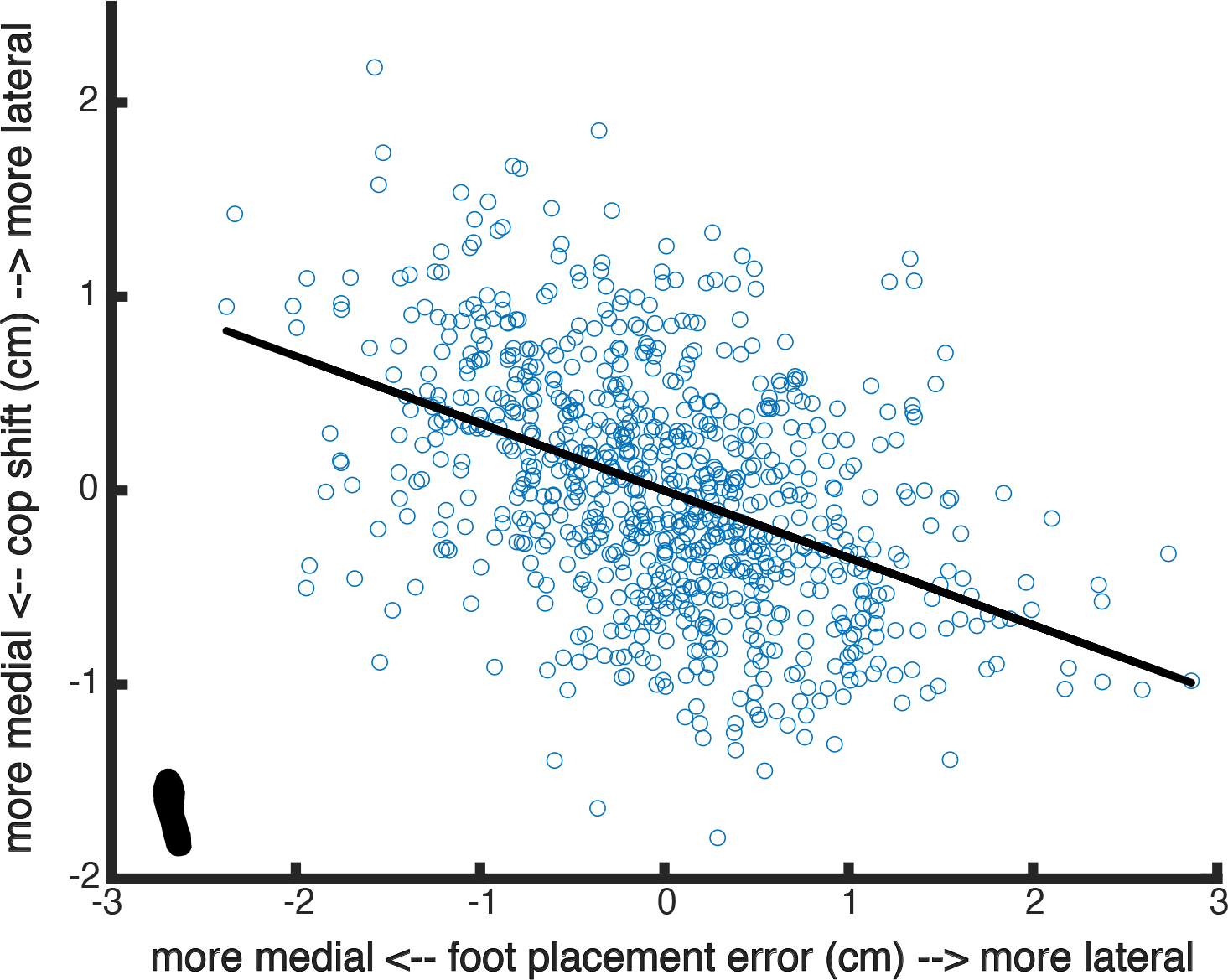
Scatter plot for Model 2 as fitted for subject 18. The slope of the relationship between foot placement error and the subsequent CoP shift is negative.

External stabilization improved gait stability, decreased step width, and diminished the degree of foot placement control (*p<0.001* for all, Figure 3).

**Figure 3.**
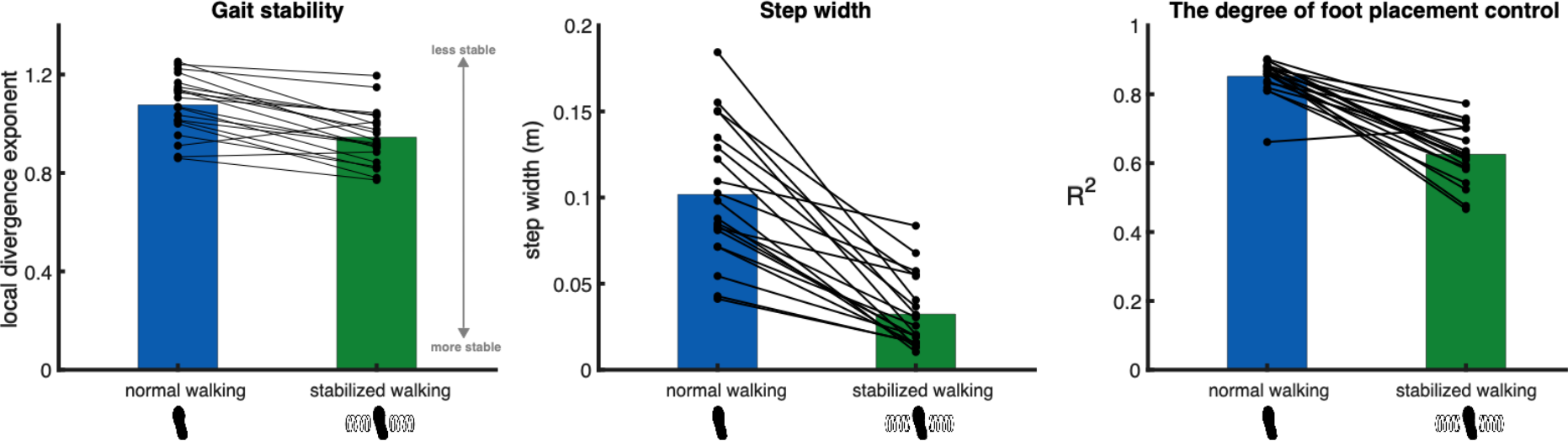
The effect of external lateral stabilization on gait stability (left panel), step width (middle panel) and the degree of foot placement control (right panel). The black dots represent the individual data points.

External stabilization significantly decreased the step-by-step ankle moment control contribution (i.e. absolute explained variance, Model 2; *p<0.05,* Figure 4).

**Figure 4.**
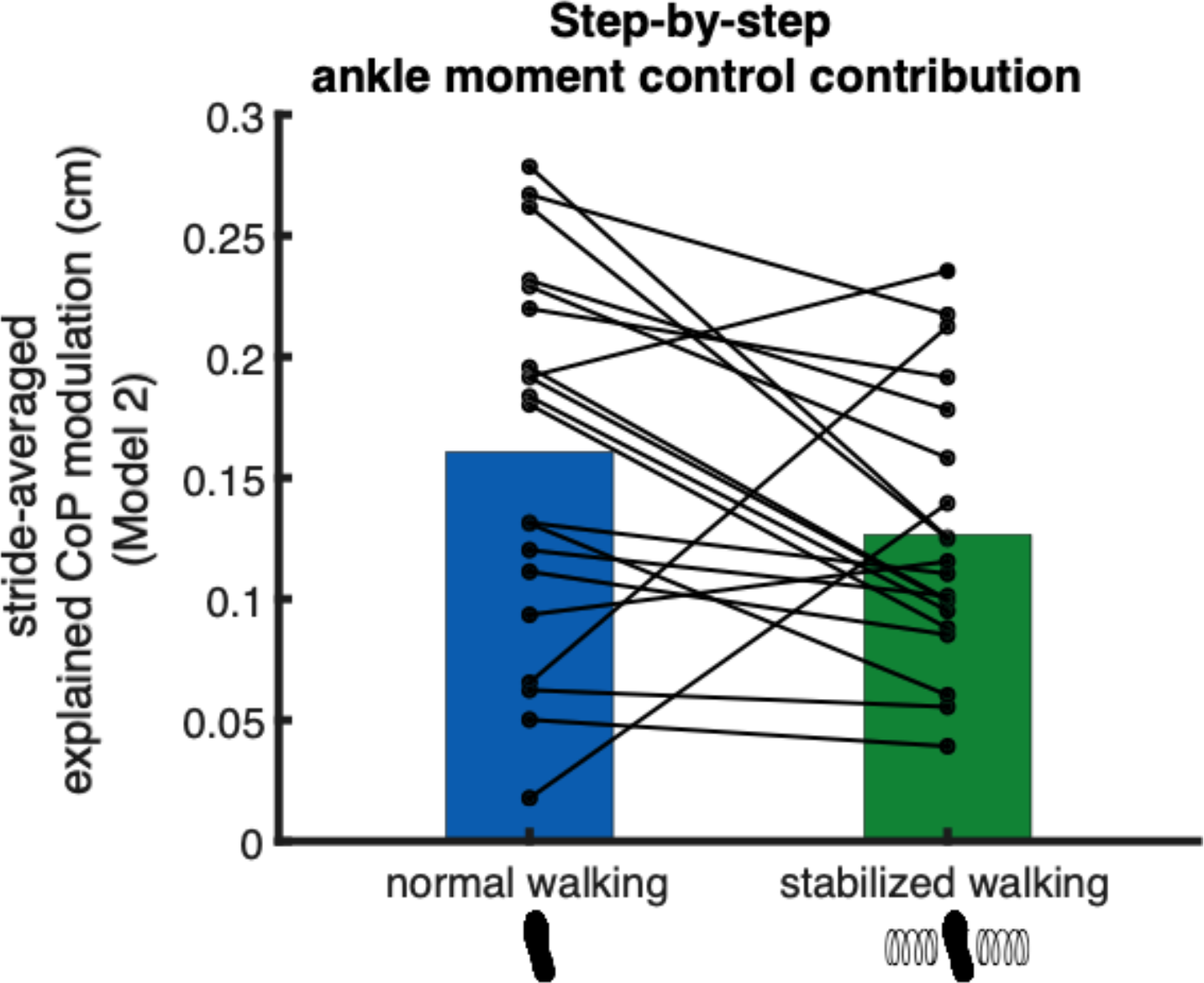
Step-by-step ankle moment control contribution (absolute explained variance, Model 2). The black dots represent the individual data points. Note that in this figure we depicted the square-root of the stride-averaged explained variance (rather than the absolute explained variance tested in the statistics), for a more intuitive interpretation of its magnitude.

External stabilization did not affect the average CoP shift (*p=0.336*, Figure 5).

**Figure 5.**
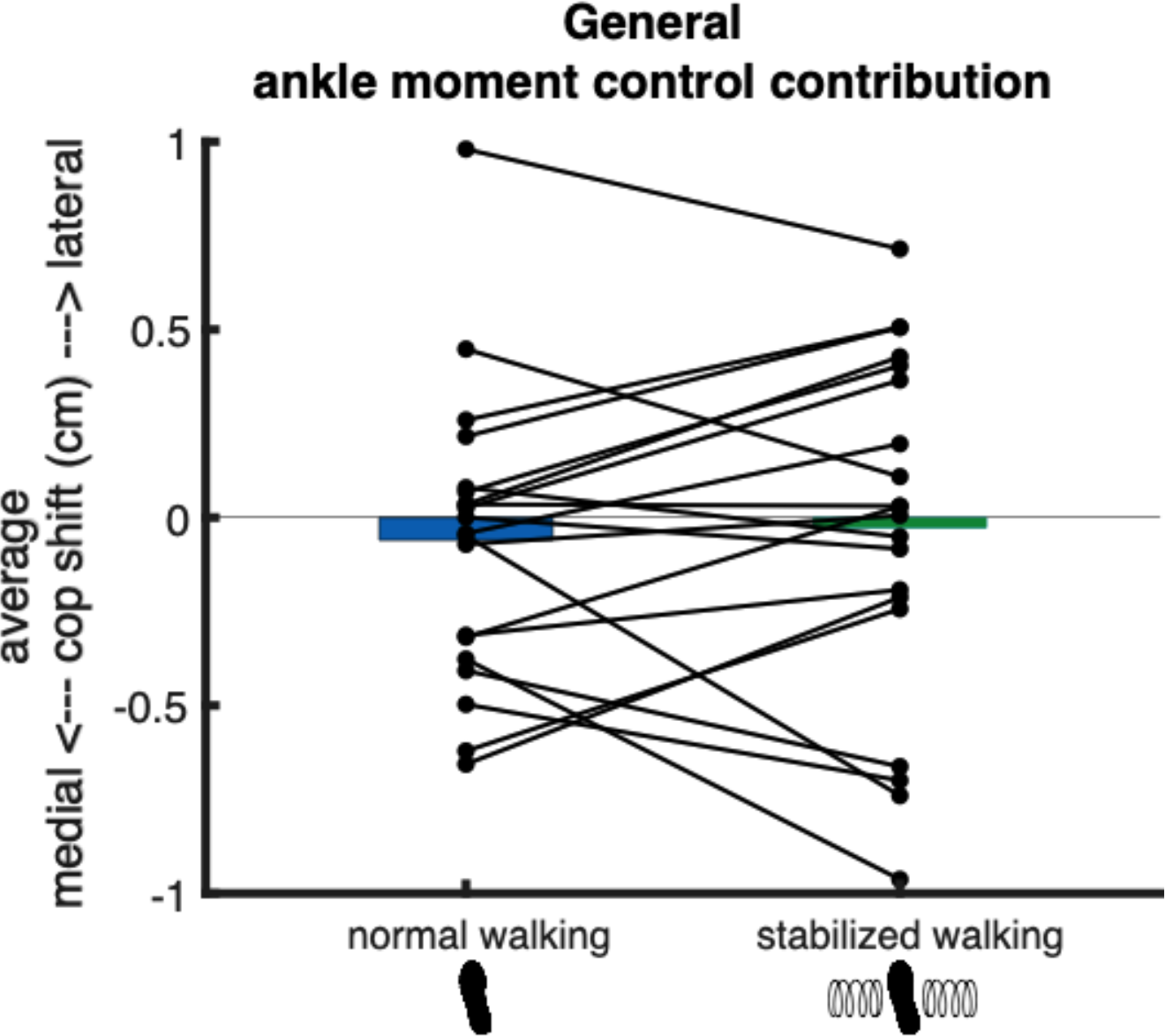
General ankle moment control contribution (average CoP shift). The black dots represent the individual data points.

External stabilization did not affect the foot placement error (*p=0.695*, Figure 6).

**Figure 6.**
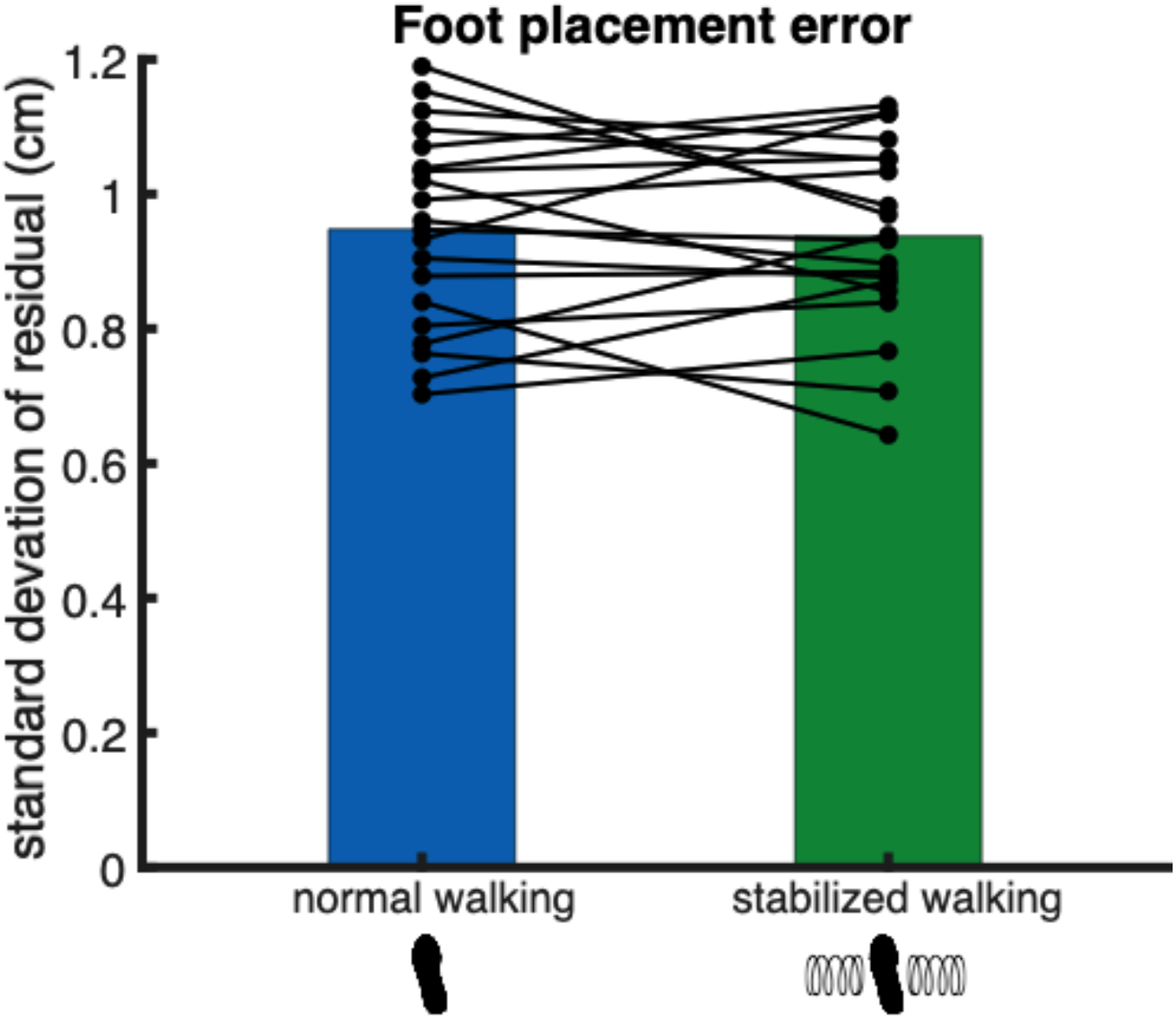
Foot placement error (standard deviation of ε_fp_). The black dots represent the individual data points.

## Discussion

On the premises that external stabilization decreases the need to actively control gait stability, we studied the role of ankle moment control in mediolateral gait stabilization. During steadystate walking, it is known that foot placement control provides a large contribution to mediolateral gait stability (Bruijn and van Dieën, 2018; Hof, 2007; Mahaki et al., 2019). A high degree of foot placement control is maintained (van Leeuwen et al., 2020; Wang and Srinivasan, 2014), ensuring appropriate coordination between the base of support and the center-of-mass. After the foot has been placed, ankle moment control can correct for errors in foot placement (Hof et al., 2007). Due to the finite width of the foot, the contribution of ankle moment control is limited. Still, the current findings are consistent with previous reports that ankle moment control partially corrects for foot placement errors (Hof et al., 2007; van Leeuwen et al., 2021). In addition, a reduction in the contribution of step-by-step ankle moment control when walking with external lateral stabilization provides evidence that ankle moment control subserves mediolateral steady-state gait stability.

### Steady-state gait stability

Not only in perturbation studies (Hof and Duysens, 2018), but also during steady-state walking, ankle moment control appears to be used to control gait stability (Fettrow et al., 2019; Hof et al., 2007; van Leeuwen et al., 2021). One way to assess whether the execution of a mechanism contributes to gait stability during steady-state walking, is by comparing the mechanism during normal walking and during stabilized walking (Mahaki et al., 2019). By taking over (part of) stability control externally, participants can loosen their control. During normal walking, participants maintain a degree of foot placement control to stabilize steady-state walking (Wang and Srinivasan, 2014). With external stabilization, participants decease this degree of step-by-step foot placement control (Bruijn et al., 2015; Mahaki et al., 2019). Moreover, when walking with lateral stabilization, participants walked with narrower steps on average, whilst walking more stable (Bruijn et al., 2015; Mahaki et al., 2019). As expected, we found similar results in the present study.

### Ankle moment control contribution

We also found that the contribution of step-by-step ankle moment control was reduced during walking with external lateral stabilization. The standard deviation of the foot placement error did not significantly differ between normal and stabilized walking. So, the absolute explained variance of the ankle moment control model (Model 2) did not decrease due to a smaller variance of the foot placement error. Rather, the reduction in ankle moment control contribution reflects a lack of foot placement error correction, due to a reduced need for active stabilization.

Unlike foot placement control, ankle moment control does not seem to have a large overall contribution to gait stability. Whilst average step width decreases when externally stabilized, the average CoP shift remained unchanged. Thus, during normal steady-state walking, the main contribution of ankle moment control appears to be in correcting step-by-step variations, rather than in realizing an average CoP shift across steps.

### Relevance

Evidence that ankle moment control is indeed used to stabilize gait in healthy individuals, may help improve stability in population groups with impaired ankle moment control. Older adults, for example, demonstrate diminished ankle moment control during perturbed walking (Afschrift et al., 2019). In addition, during steady-state walking, older adults walk with a diminished degree of foot placement control compared to young adults (Arvin et al., 2018). Their increased step width as a compensatory mechanism (Arvin et al., 2018), may reflect insufficient correction for steady-state foot placement errors through ankle moment control. Training ankle moment control may therefore benefit gait stability, and could potentially allow older adults to walk with a more energetically efficient step width (Donelan et al., 2001). In young adults, augmented proprioceptive feedback worked to enhance the degree of foot placement control (Knapp et al., 2021). Perhaps a similar paradigm can be applied for ankle moment control, to improve modulation following foot placement errors. In any case, the present study shows that step-by-step ankle moment control provides a contribution to mediolateral gait stability.

## Acknowledgements

The authors like to thank the participants and everyone who helped in the data collection. Special thanks to lab manager Mohammadreza Mahaki for instructing us on the external lateral stabilization frame. A very special thanks to Lucas Polman and Bob Goedbloed who, as talented students, ensured smooth conduction of the experiment. Sjoerd Bruijn and Moira van Leeuwen were funded by the Dutch Research Council (016.Vidi.178.014), https://www.nwo.nl/en/.

## Notes

https://doi.org/10.5281/zenodo.6994887

### Competing Interest Statement

The authors have declared no competing interest.

### Summary of Updates

In the previous version, some figure numbers in the text did not match the actual figures. We have also added the link to Zenodo where we have published our data analysis for this paper.

